# Whole-body metabolic modelling predicts isoleucine dependency of SARS-CoV-2 replication

**DOI:** 10.1101/2022.04.13.488249

**Authors:** Ines Thiele, Ronan M.T. Fleming

**Affiliations:** School of Medicine, National University of Galway, Galway, Ireland; Ryan Institute, National University of Galway, Galway, Ireland; Division of Microbiology, National University of Galway, Galway, Ireland; APC Microbiome Ireland, Cork, Ireland; Leiden Academic Centre for Drug Research, Leiden University, Leiden, The Netherlands

## Abstract

We aimed at investigating host-virus co-metabolism during SARS-CoV-2 infection. Therefore, we extended comprehensive sex-specific, whole-body organ resolved models of human metabolism with the necessary reactions to replicate SARS-CoV-2 in the lung as well as selected peripheral organs. Using this comprehensive host-virus model, we obtained the following key results: 1. The predicted maximal possible virus shedding rate was limited by isoleucine availability. 2. The supported initial viral load depended on the increase in CD4+ T-cells, consistent with the literature. 3. During viral infection, the whole-body metabolism changed including the blood metabolome, which agreed well with metabolomic studies from COVID-19 patients and healthy controls. 4. The virus shedding rate could be reduced by either inhibition of the guanylate kinase 1 or availability of amino acids, e.g., in the diet. 5. The virus variants achieved differed in their maximal possible virus shedding rates, which could be inversely linked to isoleucine occurrences in the sequences. Taken together, this study presents the metabolic crosstalk between host and virus and emphasis the role of amino acid metabolism during SARS-CoV-2 infection, in particular of isoleucine. As such, it provides an example of how computational modelling can complement more canonical approaches to gain insight into host-virus crosstalk and to identify potential therapeutic strategies.

## Introduction

Covid-19 is an infection of the respiratory tract caused by the severe acute respiratory syndrome corona virus-2 (SARS-CoV-2) [1]. It is characterised by a wide range of symptoms, including cough, fever, diarrhoea, and shortness of breath, depending on the disease severity [1]. The severity of Covid-19 varies between infected individuals, ranging from asymptomatic to critical, severe pneumonia, with multiple organ failure as a leading cause of death [1]. Several susceptibility factors and pre-dispositions, such as age, sex, and co-morbidities, have been linked to disease severity and outcome [1-3]. In addition to affecting the respiratory system, SARS-CoV-2 affects other organs [4], which has been mainly attributed to the broad expression of the receptors (hACE2) to which SARS-CoV-2 binds [5]. Accordingly, SARS-CoV-2 has been found in various organs beyond the lung, such as the liver [6], adipose tissue [7, 8], and small intestine [9-11]. The brain can also be affected leading to clinical phenotypes, such as cognitive impairment [12, 13]. Hence, Covid-19 has been recognised as a multisystem disease involving numerous organs [4].

Viral infections are known to hijack the metabolism of the infected cells [14], as they require the host’s cell metabolic machinery to produce the viral particles [5]. Consequently, the viral infection could lead to substantial alterations of cellular metabolism in the SARS-CoV-2 infected organs. Hence, the question is whether the metabolic changes do not only occur on a cellular or organ level but rather involve metabolic reprogramming on a whole-body scale, which could underlie multi-organ failure.

The disentanglement of the complex host-virus interactions can be aided by computational modelling. One suitable approach is the constraint-based reconstruction and analyses approach (COBRA) [15], in which the metabolic network of a target organism is constructed based on genomic, biochemical, physiological, and phenotypic data [16]. The metabolic reconstruction can then be used to interrogate emergent, functional properties [15]. The COBRA approach has been successfully applied to a variety of biomedical questions [17, 18], including host-pathogen interactions [19]. One of the advantages of metabolic reconstructions is that they can be tailored to a particular question or condition through the application of constraints [15], which could, for example, be dietary availability [20-22], gene defects [23, 24], or omics data [25-27]. Hence, one metabolic reconstruction can give rise to many condition-specific models. Moreover, the metabolic models of different organisms can be combined into a “superorganism” metabolic model, which allows the investigation of the metabolic interactions between the modelled organisms. For example, host-pathogen models have been investigated to understand the interplay between the host and pathogen metabolism and to predict potential drug targets. A few host cell – Covid-19 models have been generated to provide insights into cellular reprogramming and to propose anti-viral drug targets [28-30].

We have recently generated sex-specific, whole-body organ-resolved reconstructions of human metabolism (WBMs) [21], which account for 28 and 32 organs in the male and female reconstruction, respectively, as well as 13 biofluids (Figure 1A). The WBMs capture more than 80,000 reactions and have been constrained using metabolomic and physiological data to correspond to a reference man and woman [21, 31]. Thus, the WBMs are ideally suited to infect the lung with SARS-CoV-2 and investigate host-virus metabolic interactions on a whole-body level (Figure 1B). Importantly, WBM-virus modelling represents a complementary approach to observational and interventional clinical studies, providing novel insights on the systemic consequences of COVID-19 infection and replication, which would be otherwise not possible.

**Figure 1:**
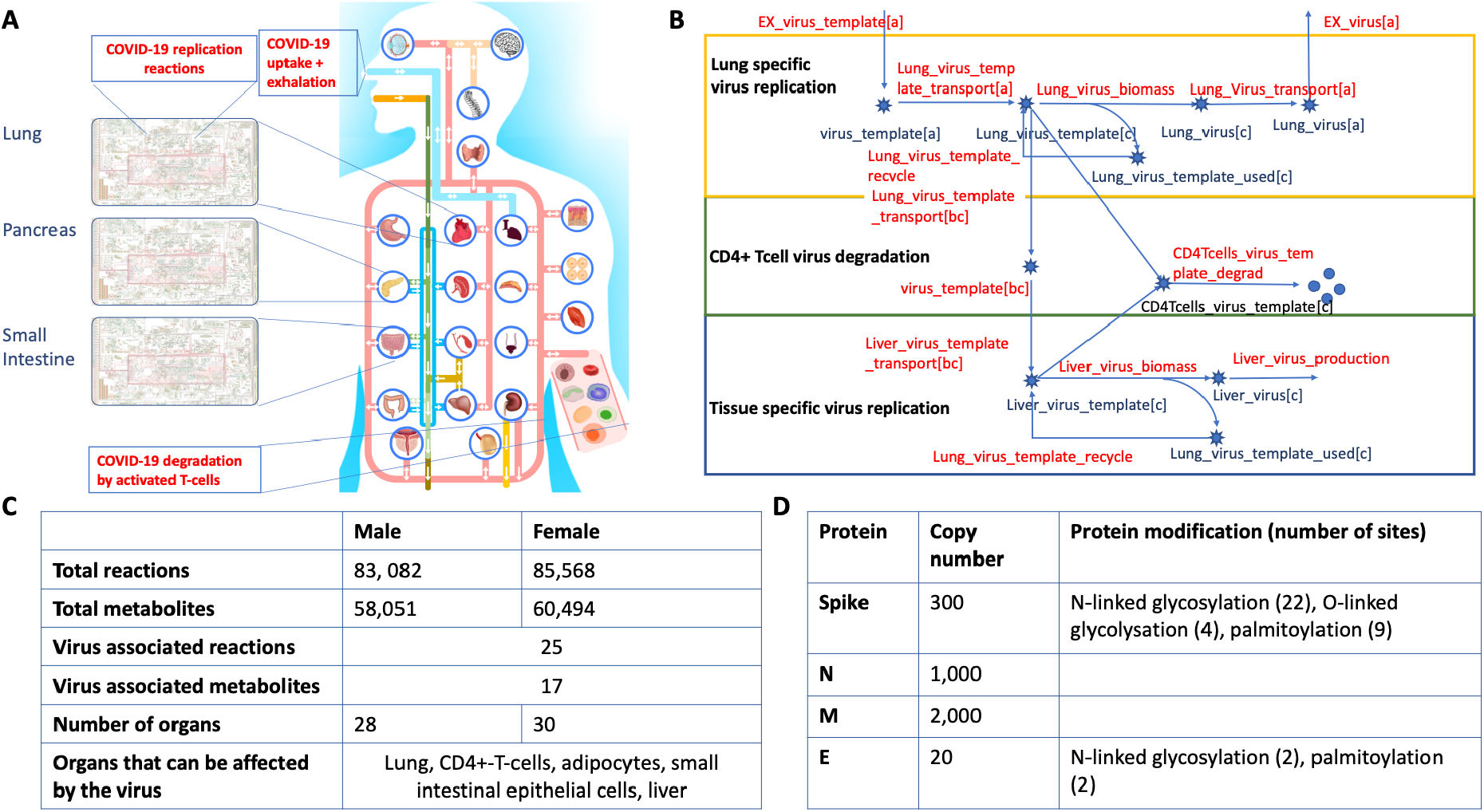
Overview of the sex-specific host-virus metabolic whole-body models. **A**. Schematic overview of the host-virus model. **B**. Schematic overview of the virus metabolic reactions added to the sex-specific, organ-resolved whole-body human metabolic models. **B**. Statistics on reaction and metabolite content of the host-virus models. **D**. Structural viral proteins, their copy numbers used for modelling the viral infection, and their protein modifications (see Methods for more details).

## Methods

### Whole-body metabolic reconstructions

We used the sex-specific organ-resolved whole-body reconstructions of human metabolism, WBMs (v1.03), with default constraints [21] (Figure 1A). Briefly, the male reconstruction consists of 81,094 reactions and 56,452 metabolites distributed across 28 organs. The female reconstruction consists of 83,521 reactions and 58,851 metabolites across 30 organs. Both reconstructions account for 13 biofluid compartments, including a blood compartment, which supplies the different organs with metabolites. The whole-body reconstructions were built from the generic human metabolic reconstruction, Recon 3D [32], using organ-specific literature, proteomic, and metabolomic data [21]. Both WBMs have been converted into personalised, condition-specific computational models using physiological and phenomenological data from a reference male and female [31] as well as constraints derived from blood, cerebrospinal fluid (CSF), and urine metabolite concentration data retrieved from the human metabolome database (HMDB) [33]. Here, we added reactions involved in *N*-linked glycan metabolism from Recon 3 [32] to various organs as they were missing from the WBM reconstructions and were required for the *N*-linked glycosylation of the spike and envelope proteins of COVID-19 (Table S1). Furthermore, additional metabolites and reactions have been added to allow better mapping of metabolomics data (Table S1). The final male reconstruction contained 83,082 reactions, 58,501 metabolites (Figure 1C), and 105,479 coupling constraints [21, 34]. The final female reconstruction contained 85,568 reactions, 60,494 metabolites, and 109,294 coupling constraints (Table S1).

### Metabolic reconstruction of virus replication

The human angiotensin-converting enzyme 2 (hACE2) serves as a receptor for SARS-CoV-2 enabling virus endocytosis via virus spike protein interaction [1]. The receptor has been found in numerous organs and tissues based on antibody staining and RNA expression, including the lung, gastrointestinal tract, liver, and adipose tissue [7, 8]. Additional evidence supports virus replication in the intestine [9-11] and the liver [6]. Upon endocytosis, the virus releases its genome, which consists of a positive-stranded RNA (+ssRNA) [35]. The virus genome contains 12 open reading frames, encoding for at least 5 accessory, 15-16 non-structural, and four structural proteins [35]. The virus is then assembled intracellularly in a dynamic compartment between the endoplasmic reticulum and the Golgi (ERGIC) [36], where the virus also gets its membrane envelop [5]. The nascent virus particle is released from the host cell via exocytosis.

The ssRNA strand of SARS-CoV-2 is translated by the host cell translation machinery, leading to the production of virus non-structural proteins that are replicating the RNA strand, resulting in the reverse (negative) RNA strand, which serves as a template for the translation of the structural proteins and for the positive strand, which will be incorporated in the nascent virus particle [35]. SARS-CoV-2 has four envelope proteins, being spike (S), envelop (E), membrane (M), and nucleoprotein N [35]. At the time of the model formulation (May 2020), the copy numbers were unknown for SARS-CoV-2. However, SARV-CoV has an estimated structural protein stoichiometry of 1S3:16M:4N to 1S3:25M:4N proteins and additional N proteins throughout the virion core [37]. Moreover, it has been reported that an averaged sized (i.e., with a diameter of 120 nm [38]) SARS-CoV virus particle contains about ∼50 to 100 spike trimers and ∼200 – 400 copies of N [37]. For the E protein, about 15-30 copies have been estimated to be present in the transmissible gastroenteritis coronavirus, as no information was available for SARS-CoV-2 or SARS-CoV [39]. Additionally, the SARS-CoV-2 virus particle contains a small but unknown quantity of accessory proteins [40]. The structural proteins are heavily modified, including N- and O-linked glycosylation of S and E (Figure 1D). In SARS-CoV, S has 22 N-linked glycosylation sites per monomer [41] and E has two potential N-linked glycosylation sites [42]. Furthermore, the cytoplasmic C-terminal end of the SARS-CoV S and E proteins are palmitoylated through the addition of palmitate to a cysteine residue via a thioester linkage [43] carried out by host proteins [44]. The S protein has nine potentially palmitoylated cytoplasmic cysteine residues, whereas E has two to three potentially palmitoylated cytoplasmic cysteine residues [43].

Based on this information, we formulated the biomass reaction, which represents the virus replication, for the SARS-CoV-2 virus using SARS-CoV-2-specific information where available or substituted it with related coronavirus data otherwise. For the biomass formulation, we followed the workflow provided elsewhere [16, 28, 45]. First, we obtained the COVID-19 genome and protein sequence from NCBI (NC_045512.2, May 2020). The fasta file contained 13 annotated (poly)proteins. The nucleotides of the genome sequence were counted for the negative and the reverse strand. The amino acids were counted for each (poly)protein and multiplied by the respective copy numbers (Figure 1D). We assumed that i) only one negative ssRNA strand, ii) 300 S proteins [37, 46], 1000 N proteins [37, 46], 2000 M proteins [37, 46], and 20 E proteins [39], iii) one copy of each of the remaining, non-structural are present in each virus particle. The precursor requirements for the *N*-linked glycosylation were calculated assuming that all 22 potential glycosylation sites per S monomer [41] and two per E protein are glycosylated [42] and using the aforementioned protein copy numbers. All four *O*-linked glycosylation sites in the S protein were assumed to be glycosylated [47]. All palmitoylation sites were assumed to be palmitoylated (nine for S and two for E) [43]. The molecular weight of the virus particle was calculated accordingly. The energy cost (adenosine triphosphate (ATP) requirement) for nucleotide sequence polymerization was assumed to require the hydrolysis of one ATP to adenosine diphosphate (ADP) and orthophosphate (Pi). The formation of a peptide bond was assumed to require four ATP. The virus replication reaction was added to the lung, liver, adipocytes, and small intestinal enterocytes of the WBMs (Figure 1B).

Additionally, we added the following reactions to the WBMs to complete the virus metabolic reconstruction. A virus uptake reaction (EX_virus_template[a]), representing the inhalation of the virus to both WBM reconstructions, and a virus shedding reaction (EX_virus[a]). An import and export reaction of the virus template and the virus particle, respectively, to and from the lung (Figure 1B). A virus accumulation reaction for the liver (Liver_virus_production), the adipocyte tissue (Adipocytes_virus_production), and the small intestinal epithelium (sIEC _virus_production) (Figure 1). The virus template can be transported from the lung into the blood circulation (Lung_virus_template_transport[bc]) and then be taken up by the liver (Liver_virus_template_transport[bc]) and the adipocytes (Adipocyte_virus_template_transport[bc]). The small intestinal epithelium can take up the virus template from the air representing that the virus can enter the gastrointestinal tract (lumen, [lu]) via the mouth and/or nasal cavity (sIEC_virus_template_transport[lu]). The viral replication in the peripheral organs does not contribute to the viral shedding flux [EX_virus[a]) representing newly produced viruses leaving the host via the airways.

To account for the host immune response, we added the uptake and degradation of the virus template to the reconstructions of the CD4+-T-cells present in the WBMs (CD4Tcells_virus_template_degrad), which accounts only for the breakdown of the ssRNA into its constituents but not of the viral shell (i.e., the amino acids). We did not represent the release of cytotoxins by the CD4+-T-cells and the host cell death upon cytotoxin release. The resulting WBM models were deemed WBM-SARS-CoV-2. In recovering COVID-19 patients, increased CD8+-T-cell and increased CD4+-T-cell levels have been found [48]. Accordingly, we created a female and male model that represents the increase in T-cells, deemed WBM-SARS-CoV-2-CD4+. The CD4+-T-cells were chosen as the WBMs do not account for CD8+T-cells. All virus-related reactions can be found in Table S2.

### Initial viral load

We enforced, with coupling constraints [21, 34], that the increases in replication reaction flux (i.e., of the virus biomass reaction) were linked in the models to higher virus template uptake flux (e.g., EX_virus_template[a]). This is the case for all four organs. The upper, or maximal, coupling coefficient was set to 2,000, in accordance with the estimate that about 1,000 viruses are produced from one virus per 10 hours [46]. In addition, we also enforced that when a virus template is taken up that the lung has to produce at least as many viruses as there were taken up. This was not the case for the other tissues.

### Diets

All simulations have been carried out using an average European diet [49] if not specified differently. Diet formulations were taken from the Virtual Metabolic Human (vmh.life) database [49]. Each diet formulation defines the uptake rates for the different dietary constituents (Table S7). The diets do not differ in the metabolites (or constituents) but rather in their overall contribution to the diet. The list of constituents is likely to be incomplete as only up to 132 metabolites are regularly measured and reported per food stuff and hence, were thus included in the diet database of the vmh.life [49].

### COBRA modelling and flux balance analysis

The sex-specific WBM-SARS-CoV-2 and WBM-SARS-CoV-2-CD4+ models were mathematical representations of the host and virus metabolic transformation and transport reactions that were parameterised as described above. The COBRA approach assumes the modelled system to be at a steady state meaning that the change in metabolite concentration (dx) over time (dt) is zero. The underlying system of linear equations can be efficiently solved using linear programming. Generally, an objective function is either maximised or minimised subject to mass-balance constraints as well as other imposed constraints (e.g., nutrient uptake). This approach is called flux balance analysis [50]. If not stated differently, we used the virus shedding reaction (‘EX_virus[a]’) as an objective function and maximised the flux through this reaction. The resulting solution contains a flux value for each reaction in the WBM-SARS-CoV-2 or WBM-SARS-CoV-2-CD4+ models. However, due to the degenerative nature of the underlying linear programming problem, i.e., we have more reactions (variables) than mass balances (equations), the solution vector is non-unique (but the objective value is). To obtain a reproducible, unique solution (out of the set of infinite possible solutions), one can minimise the Euclidean norm of the solution vectors upon maximisation of the objective function. The resulting solution vector is assumed to be closest to a thermodynamically feasible flux distribution. These flux distributions were used for the overlay onto the human metabolic map (ReconMap [51]) hosted at the VMH [49] using the corresponding functions in the COBRA Toolbox [52]. To determine metabolites, which limit higher values through the objective function, here the virus shedding reaction, we investigated the shadow prices, which are dual to the linear programming problem [15].

### Prediction of the blood metabolome

The *in silico* blood metabolome was determined by adding to each model individually for each of the 1,033 metabolites in the blood compartment ([bc]) a demand reaction (e.g., for D-glucose: DM_glc_D[bc]: 1 glc_D[bc] → Ø). These artificial reactions break the steady-state assumption and allow for the accumulation of the respective metabolite in the blood compartment. We then maximised each demand reaction individually in the sex-specific version of the healthy WBMs, WBM-SARS-CoV-2, and WBM-SARS-CoV-2-CD4+ models. We compared the flux through the respective demand reactions between the three models for each sex. A metabolite was considered increased in the blood compartment of the maximal possible flux was higher in the SARS-CoV-2 infected models compared to the healthy WBM model of the same sex and decreased if the maximal possible flux was smaller. All objective flux values less than 1e^-6^ were considered to be zero. The comparison with published plasma metabolomic data was done by mapping the reported metabolite names into the namespace of the virtual metabolic human database [49], which is also used for the WBMs (Table S5, S6). To test whether the prediction accuracy was statistically significant, we chose 1,000,000 random sets of 103 metabolites and their predictions from the 1,033 blood metabolites and compared them with the measured changes.

### Variant and sequence-specific modelling

We downloaded the COVID-19 sequences for the different variants from GISAID (https://www.gisaid.org/, between June 2021 and February 2022). To ensure that we only obtained high-quality sequences, we required the sequences to be complete, to have high coverage, with patient status, and to have complete collection dates. All submissions to GISAIDS must have been done after 01.01.2020, and the collection date must also have been on or after that date. Where necessary, we specified dates and/or geographical locations to reduce the number of downloaded sequences. We then used Diamond [53] with default parameters to perform blastp for each downloaded sequence against the reference strain (NC_045512.2). For further computational analysis, only those variants/sequences were used, for which all 13 (poly)proteins could be identified and contained no duplicates.

For each sequence, we adapted the female WBM-SARS-CoV-2 model by formulating a sequence-specific virus_biomass reaction assuming that the copy number of each protein remained the same (Figure 1D) but adjusting based on the frequency of nucleosides and amino acids based on the sequence. We also assumed that the protein modifications (Figure 1D) were not affected by mutations. Each of these variant sequences resulted in a sequence-specific model. We maximised the virus shedding reaction for each sequence-specific model. To compare the amino acid frequency of each model, we added the occurrence of each amino acid per protein multiplied by the copy number of the protein.

## Code availability

All simulations were carried out using the COBRA Toolbox v3.0 [52] and the PCSM toolbox [21] using Matlab 2020 (Mathworks, Inc) as simulation environment and Ilog clpex (IBM, Inc) as linear programming solver. The code is available in the GitHub repository of the COBRA Toolbox: https://github.com/opencobra/COBRA.papers (after publication).

## Results

### Generation of sex-specific host-virus metabolic whole-body models

To model SARS-CoV-2 viral infection, we expanded the comprehensive, organ-resolved, sex-specific whole-body models of human metabolism (WBM) [21] with SARS-CoV-2 specific reactions (Figure 1B). These reactions were formulated based on available data on SARS-CoV-2 and related coronaviruses (Method section). Briefly, the virus is taken up from the air (EX_virus_template[a]) and then by the lung where the virus replicates (viral biomass reaction, VBR). The resulting virus particles leave the lung and are released into the air (EX_virus[a], Figure 1B). The VBR was formulated such that it accounts for all known viral biomass precursors, being 1. the nucleotides for the single-strand RNA (ssRNA), 2. the amino acids for the structural and non-structural viral proteins encoded by the ssRNA, 3. the *N*-linked and *O*-linked glycans present on the spike and the envelope proteins, and 4. the palmityl-CoA required for the palmitoylation of the S and E proteins (Figure 1D). The structural protein copy numbers were retrieved from the literature (see Method section for details). The virus particle can be degraded by the CD4+ T-cells in the WBMs. We do not represent the death of lung cells and non-metabolic inflammation and immune responses. However, the WBMs contain reactions for the metabolism of immuno-metabolites (e.g., eicosanoids) and thus may capture potential changes along those pathways. Furthermore, the setup allows the virus to replicate in the liver, adipocytes, and the small intestine, consistent with reports of high expression of the human ACE2 receptor, to which the virus binds [7, 8]. In total, 25 virus-specific reactions were formulated and included in the WBMs (SI Table S2), yielding the WBM-SARS-COV-2 models consisting of 83,082 metabolic reactions for 28 organs for the male model and 85,568 reactions for 30 organs for the female model (Figure 1C). The WBM-SARS-COV-2 models were constrained based on the physiological parameters of a reference man and a reference woman (e.g., weight, height, organ contributions to the whole-body weight, blood perfusion rates of the different organs) [21]. No sex-specific or personalised constraints were placed on the virus reactions, as such data were not available. However, we limited the ratio of the virus uptake reaction (e.g., EX_virus_template[a]) to the virus biomass reaction (e.g., Lung_virus_production) to be maximally 2,000 (per day and person), representing the estimated burst size of 1,000 per 10 hours [46]. Furthermore, the dietary uptake constraints were set to correspond to an average European diet [49], if not specified differently. Taken together, we generated computational metabolic models of viral infection of the human host, which have been tailored using condition-, sex-specific constraints.

### Modelling COVID-19 infection

First, we investigated the consequences of virus replicating in the lung. In the male and the female WBM-SARS-COV-2 models, using flux balance analysis [50], the maximally possible flux through the virus shedding reaction (EX_virus[a]) was 33.0254 U (mmol/day/person) from 1 U inhaled virus (EX_virus_template[a]) (Table S3). The predictions are not quantitative and the viral uptake of on mmol/day/person cannot directly be correlated with the viral load reported in individuals [54]. Notably, in both WBM-SARS-COV-2 models, the uptake of the essential amino acid isoleucine by the lung from the blood circulation was limiting the maximally possible flux through the virus shedding flux reaction. To test whether isoleucine is indeed rate limiting, we relaxed the upper bound on the lung isoleucine uptake reaction, which was set based on the blood concentration of isoleucine in healthy individuals and the blood perfusion rate through the lung [21]. The maximal flux through the virus shedding flux reaction increased to 41.0819 U for the male model and 38.711 U for the female model when other metabolites became rate-limiting (Table S3).

The WBM-SARS-COV-2 models corresponded to a mild, not hospitalisation requiring, infection with normal amounts of CD4+ T-cells. To simulate the increased viral load reported for mild (but hospitalised) and severe COVID-19 patients (5.11 vs 6.17 log10 copies per mL, respectively, [54]), we increased the viral uptake flux to 10 U. However, no feasible solution could be obtained, which was expected as already a mild (but hospitalisation requiring) SARS-COV-2 infection led to an increase in CD8+ T-cells of approximately six times and approximately three times for CD4+ T-cells [48]. Since the WBM-SARS-COV-2 models do not account for CD8+ T-cells, we decided to use a factor of ten in the subsequent simulations to approximate the combined raise of T-cells. The increase in T-cells was modelled by adjusting the coefficient corresponding to CD4+ T-cells in the whole-body biomass reaction. The resulting models, deemed WBM-SARS-COV-2-CD4+, had a the maximally possible flux through the virus shedding reaction of 33.0254 U (female and male), when the constraint on the isoleucine uptake was unchanged or to 41.0819 U for the male model and 38.711 U for the female model when the lung isoleucine was increased to 100 U (Table S3).

These results show that in the host-virus WBMs an increase in T-cells is required to deal with a higher initial viral load, consistent with our current knowledge.

### Whole-body metabolic remodelling during Covid-19 infection

To obtain an assessment of whole-body metabolic remodelling, we pursued two alternative approaches. First, we investigated the metabolic changes associated with the infection, the increase in virus load, and CD4+ T-cell availability. Therefore, we used three models for each sex, healthy WB), WBM-SARS-COV-2 model with 1 U virus uptake and normal CD4+ T-cell levels, and WBM-SARS-COV-2-CD4+ model with 10 U virus uptake and 10 times increase in CD4+ T-cells. We calculated the flux distribution that minimises the Euclidean norm, thereby approximating the closest thermodynamically feasible flux distribution for each model. When comparing the flux distribution obtained from the WBM-SARS-COV-2 with the one from the healthy WBM model, approximately 15% of the metabolic reactions changed in flux values by at least 10% for both sexes (Figure 2A, Table S4). Similar numbers in reaction flux changes were observed when comparing the WBM-SARS-COV-2-CD4+ results with the WBM-SARS-COV-2 and with the healthy WBM model results. These results indicate an overall change in metabolism due to virus infection but also between mild and severe infection, involving almost all organs (Table S4). In the female lung, 12% of the 3,467 lung reactions increased in flux, while 14,7% of reactions decreased in flux compared to the healthy female WBM (Figure 3B). We noticed that these results were sensitive to the applied constraints and the reaction content of the WBMs, as can be seen in the differences between male and female models. This problem arose as we calculated only one of an infinite number of possible flux distributions that are consistent with the applied constraints. Nonetheless, the results illustrate that the metabolism in the entire body was affected during the viral infection, consistent with our current knowledge and reports in the literature [4].

**Figure 2.**
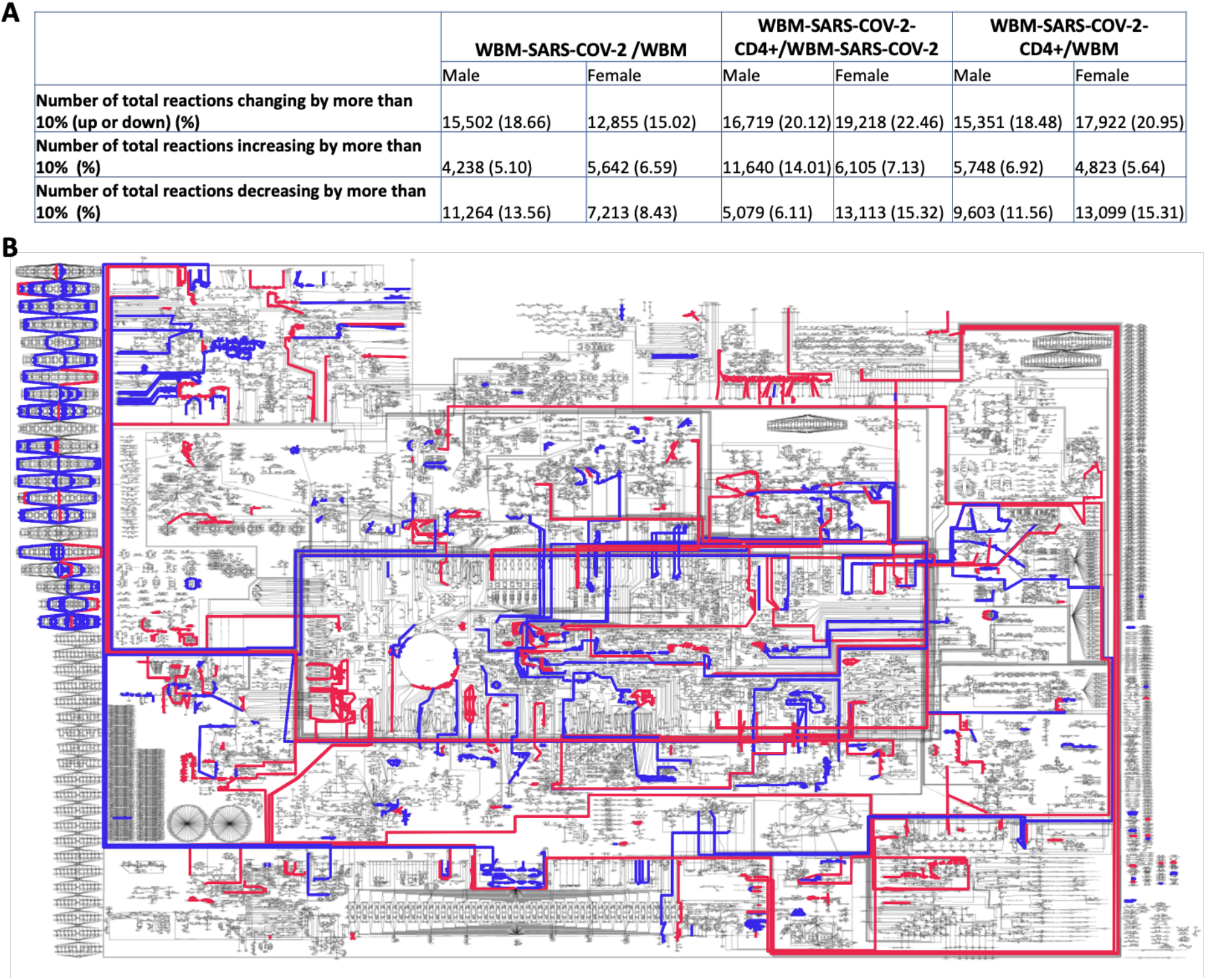
Metabolite changes occurring during mild and severe virus infection. **A**. Overall changes in reaction flux values in the WBM models in mild (WBM-SARS-COV-2) and severe (WBM-SARS-COV-2-CD4+) infection models compared with the healthy WBM models. **B**. Biochemical network visualisation of predicted metabolic changes occurring in the female lung during mild infection. Flux values that increased (red) or decreased (blue) by more than 10% when comparing the female WBM-SARS-COV-2 with the healthy WBM model.

**Figure 3:**
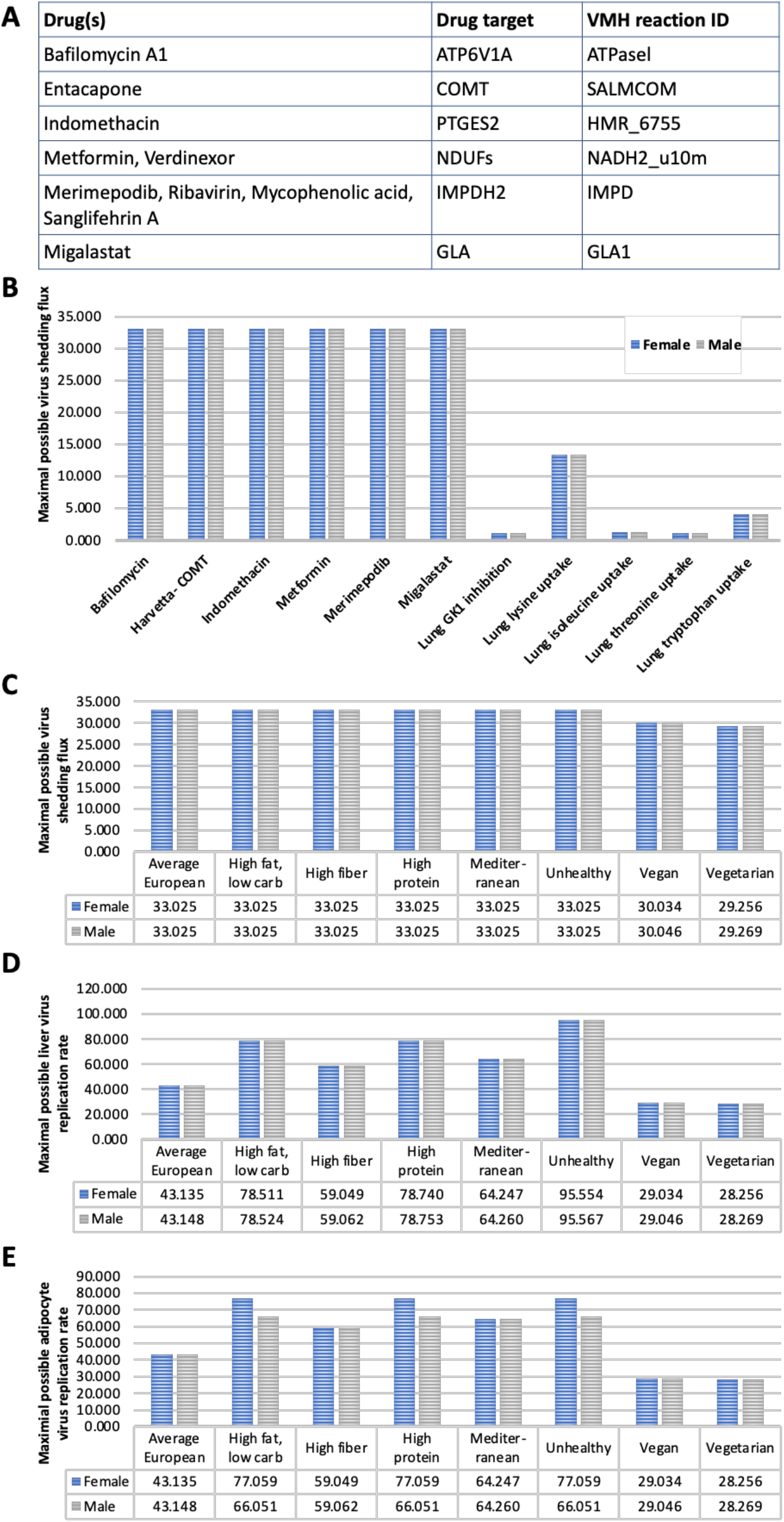
Predicted effect of drug treatment (A) and different dietary regimes on the maximally possible virus shedding rates in the different organs (B). All fluxes are given in mmol/day/person.

As a second approach, we calculated the maximally possible increase or decrease of metabolites in the blood compartment. Therefore, we added for each of the 1,033 metabolites present in the blood compartment an artificial reaction allowing for the metabolite accumulation (Methods) and maximised the flux through each reaction individually. We found that 353 out of 1,033 (34%) metabolites changed in the female WBM-SARS-COV-2 model and 359 out of 1,033 (35%) metabolites changed in the male WBM-SARS-COV-2 model (Table S5). In the female WBM-SARS-COV-2-CD4+ model, we predicted an increase of 79 and a decrease of 278 blood metabolites. Next, we compared the predicted blood metabolite changes with published metabolomic data [55], which reported 474 statistically significant changes between healthy and non-severe Covid-19 patients (n = 28 and n = 37, respectively) (Table S6). We could map 127/474 (27%) measured metabolites onto the blood metabolites in the WBMs. Of these 127, 103 had a non-zero maximal flux value through their respective demand reaction. For the female WBM-SARS-COV-2 model, the predictions agreed for in 73/103 (71%, p = 0.0006) of the cases. In six (6%) cases, we predicted a decreased metabolite level while an increase was reported. In further 24/103 (23%) of the cases, we predicted no change. The numbers were very similar for the male WBM-SARS-COV-2 model (Table S5).

Taken together, our simulation results illustrate a metabolic remodelling on a whole-body level that also affect the blood metabolome, which showed good agreement with the published data.

### Simulation of potential anti-viral drug targets

Over the past two years, various potential drug treatment strategies have been suggested. One study proposed a set of 69 FDA-approved drugs targeting 66 host proteins as potential drugs, as they had shown that these interacted with COVID-19 proteins [56]. Hence, we investigated whether any of these drugs would alter the virus shedding rate (EX_virus[a]) in our host-virus models. Only ten of the 66 host proteins were covered in the WBM models (Figure 3A). We inhibited each of these proteins by setting the upper and lower constraints on the corresponding lung reactions to zero and maximised the virus shedding flux. None of these inhibitions resulted in a decrease of the maximally possible virus shedding flux *in silico* (Figure 3B). Another study, which also used a computational model of COVID-19 and a metabolic model of human macrophages, has suggested the guanylate kinase as a drug target [28]. The guanylate kinase catalyses the reversible ATP-dependent phosphorylation of GMP to GDP. In agreement with that study, the inhibition of the corresponding reactions (VMH ID: GK1, DGK1) in the WBM-SARS-COV-2 models reduced the maximal virus production potential to about 3.3% of the original WBM-SARS-COV-2 model flux values (Figure 3B, Table S3). The complete inhibition of the associated reactions led to an infeasible model, meaning that a small residual flux through these reactions was necessary to sustain the maintenance of the WBM model. Additionally, we inhibited the lung uptake of various amino acids, motivated by the result that isoleucine was rate-limiting for the viral shedding flux (Figure 3B). The reduction of the lung uptake of isoleucine and threonine resulted in a similar reduction of maximal virus shedding flux as did the GK1 inhibition (Figure 3B). In contrast, the reduction of uptake flux in tryptophan and lysine resulted in a reduction of the maximal virus shedding flux to 12.25% and 40.3%, respectively, in both sexes. These results highlight the potential of metabolic targets for reducing the viral replication rate within the lung tissue.

### Predicted effect of diet on virus replication in various organs

The reduction in lung amino acid uptake could be achieved by, e.g., dietary changes. Hence, we investigated whether different diets may alter the maximal possible viral shedding rate. Therefore, we altered the *in silico* diet of the WBM-SARS-COV-2 and WBM-SARS-COV-2-CD4+ models and maximised the maximally possible virus shedding rate (EX_virus[a]) (Figure 3C, Table S3). In total, we used eight different diets from the Virtual Metabolic Human database (vmh.life) [49]. The lowest maximally possible virus shedding rate was predicted for the vegan and vegetarian diets (Figure 3C), which also have the lowest isoleucine content (Table S7), while all other diets resulted in the same maximal shedding flux as the average European diet. Interestingly, tryptophan became the rate-limiting amino acid in the vegan and the vegetarian diet (Table S3). Both diets have the lowest tryptophan content (Table S7). Next, we investigated whether diet may influence the virus replication rate in the other infected organs. The maximally possible virus replication fluxes were dependent on the diet in the liver and the adipocytes. In contrast, in the small intestine, the flux was limited by a glycan (VMH ID: g3m8mpdol_L), which was required for the N-linked glycosylation of the S and E proteins (Figure 1D, Table S3). The vegan and the vegetarian diet resulted in the lowest liver and adipocyte virus replication fluxes. In the liver, the highest virus replication flux was obtained for the unhealthy diet, which has been defined as a burger- and steak rich diet [49], followed by the high fat and the high protein diets (Figure 3D). The virus replication fluxes were limited by tryptophan in the liver and the adipocytes (Table S3). Sex-specific differences were obtained for the unhealthy, high fat, and high protein diets in the adipocytes (Figure 3E), which can be attributed to women having a higher percentage of body fat, which was also the case for our female WBM (Figure 3E). The results were similar for the WBM-SARS-COV-2-CD4+ models (Table S3). Taken together, these results suggest that diet and sex could influence the virus shedding flux in the different infected organs.

### Analysis of virus variants

Numerous SARS-CoV-2 variants have been identified over the past two years, some of which have been the drivers behind the different pandemic waves. In February 2021, the World Health Organisation introduced a naming and monitoring system, which lead to the classification of variants under monitoring (VUM), variants of potential interests (VOI), and variants of concerns (VOC). So far, we have used for our *in silico* investigations the original, or parental, virus genome sequence, which was released in February 2020. To investigate whether the mutations found in the different variants may have adapted not only to evade the immune response, through mutations of critical amino acids in the spike protein, but also to the host metabolism, we obtained genome sequences from GISAID for the five VOCs, two VOIs, and one VUM (classification status of December 2021, Figure 4A). Furthermore, to have more sequences that are likely to be closest to the parental virus, we also obtained sequences from Italy in February 2020, where the first wave in Europe started and that resulted in many deaths (Figure 4A). We then created a total of 12,233 WBM-SARS-COV-2 models (Methods), each specific for a virus sequence, and maximised the virus shedding reaction. Interestingly, the predicted virus replication rates varied significantly between the variants (Figure 4B). The delta variant achieved the highest shedding rate followed by B.1.640, a variant under monitoring by the WHO, which had been reported in France in December 2021 [57], and its occurrence overlapped with the omicron wave. The maximal virus shedding rate predicted for the omicron variant, which now represents the dominant variant worldwide, was lower than that of the parental strain (Figure 4B, Table S8). The variant sequenced in Italy in Spring 2020 had an average maximal virus shedding rate comparable to the parental variant but was lower than the delta variant and the B.1.640 variants. However, only 69 sequences of B.1.640 had been deposited at GISAID at the time of analysis, and of those, only 28 passed our stringent quality requirements (see method). The subvariant of the omicron variant, deemed BA.2, has overtaken the omicron variant BA.1 since January 2022. Its predicted maximal virus shedding rate BA.2 was slightly higher than that of BA.1 (Figure 4B, Table S8).

**Figure 4.**
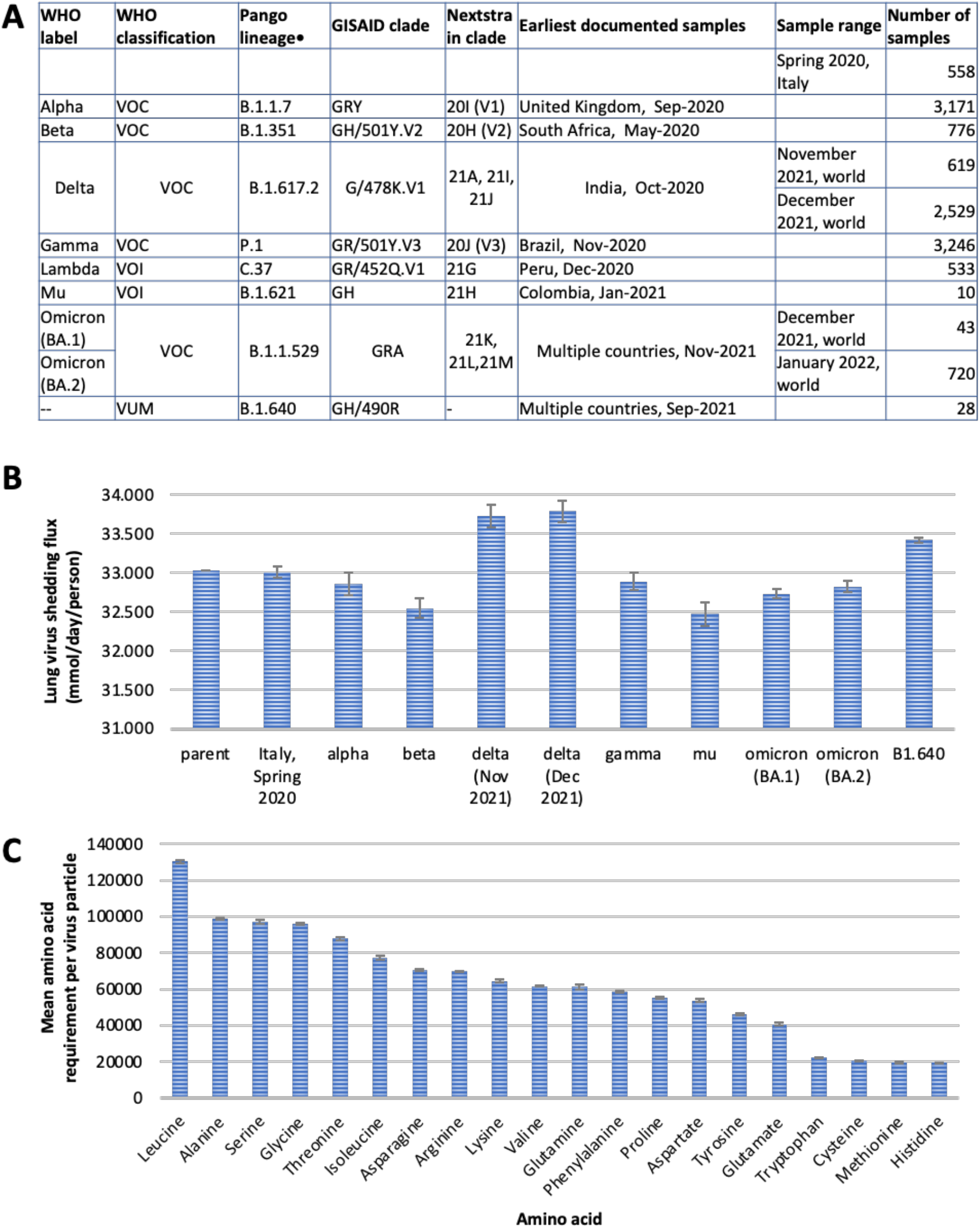
Analysis of SARS-CoV-2 variants in the context of the host-virus whole-body models. **A**. List of considered variants and their classifications. A total of 12,233 variant sequences were analysed. **B**. Predicted maximal possible virus shedding flux using the variant-specific WBM-SARS-COV-2 models. **C**. Mean amino acid requirements per particle were determined by multiplying the number of amino acids in the structural and non-structural proteins by the protein copy numbers (Figure 1D).

To better understand the variation in replication rates, we analysed the amino acid composition of COVID-19 variants (Figure 4C, Table S9). As the sequence itself is not sufficient for the amino acid requirements to create a new virus particle, we multiplied each amino acid frequency with the copy number of the viral proteins (Figure 1D). We found that the highest requirements were for leucine, alanine, glycine, serine, and threonine, while the lowest requirements were for histidine, methionine, cysteine, and tryptophan (Figure 4C). Curiously, but consistent with the simulation results using the parental variant, the predicted virus shedding flux increased linearly with decreasing isoleucine abundance in the viral proteome (R^2^=0.99, Figure 5A, Table S8). In contrast, we observed that the virus exhalation flux increased linearly with increasing threonine requirements, except for the omicron subvariants (Figure 5B, Table S8, Figure S1). The omicron threonine requirements were comparable to those of the delta variant, but its replication rate was limited by its high requirement for isoleucine.

**Figure 5.**
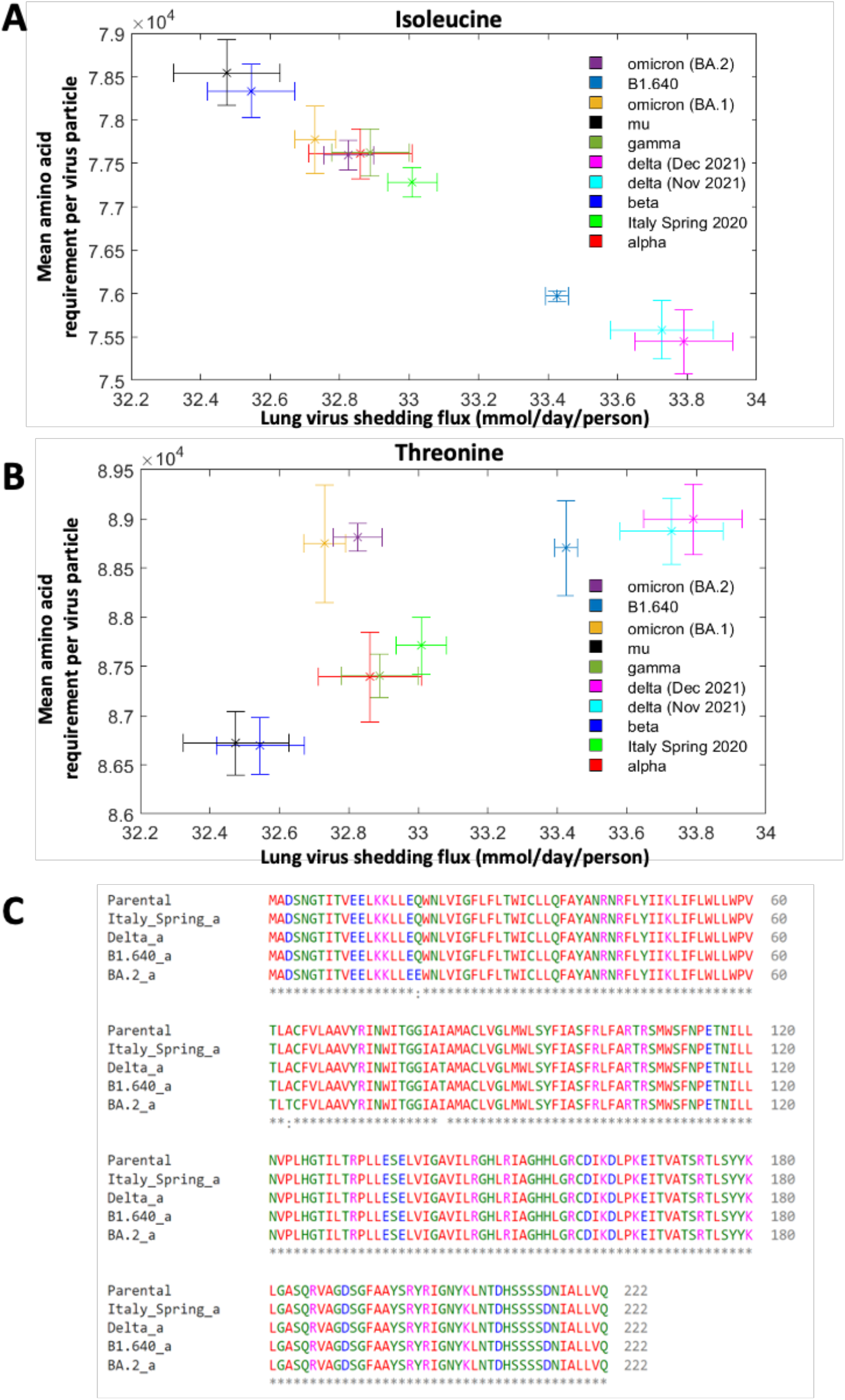
Predicted dependency of the maximal possible virus shedding flux on isoleucine and threonine requirements of the nascent virus particle. Predicted anticorrelation of the maximal possible virus shedding flux in the variant-specific female WBM-SARS-COV-2 models on isoleucine (**A**) and threonine (**B**). **C**. Sequence alignment of randomly chosen sequences of three variants and the parental virus for the M protein. See also Figure S2-4 for more examples.

The Covid-19 variants are defined by their mutations, which differ in type (i.e., nucleotide change) as well as in the affected proteins (Figure 4F, Figure S2-4). Much attention has focused on mutations in the spike protein, which is of high importance as the vaccines have been developed to enable immune system recognition of certain parts of the spike protein. However, less attention has been given to the other open reading frames. For instance, in the delta and the B.1.640 variant, one isoleucine has been substituted in the M protein with threonine when compared with the parental variant (Figure 5C). However, since one virus particle contains 2,000 M proteins, this single non-synonymous replacement led to the greatly reduced requirement for the essential amino acid isoleucine and thus, to an increased maximal possible virus shedding rate. The B.1.640 variant had one additional isoleucine in the N protein (Figure S3) reducing its maximal possible shedding flux compared to the delta variant. In contrast, the omicron variants did not contain this mutation (Figure 4F). Overall, the sequences of the delta variants were decreased in aspartate and isoleucine content, while increased in cysteine and methionine content, compared to the variant that caused the Italian wave (Table S9). Omicron, in contrast, was mostly decreased in glutamine (Figure S2) and increased in lysine (Figure S4) content. However, for all variants lung isoleucine uptake remained rate-limiting for the virus shedding flux.

Taken together, single mutation and their combinations in the structural proteins can lead to differences *in silico* virus shedding rates and may reflect potential adaptation to metabolic properties of the host.

## Discussion

We investigated host-virus co-metabolism during SARS-CoV-2 infection. Therefore, we extended the comprehensive sex-specific, whole-body organ resolved models of human metabolism with the necessary reactions to replicate SARS-CoV-2 in the lung as well as selected peripheral organs. We used the host-virus model to predict the maximal possible virus shedding rate *in silico*. To replicate within the *in silico* host cells, the virus template is transcribed and translated, and the nascent proteins may be further modified (Figure 1B) [5]. As such, the viral infection poses an additional metabolic burden on the *in silico* host and leads to the re-direction of metabolic fluxes towards the viral replication. Our predictions of large-scale metabolic changes in metabolic fluxes (Figure 2A) are thus consistent that any viral infection leads to remodelling of cellular metabolism [14, 58, 59]. However, the identification of potential bottlenecks, or rate-limiting steps, might be difficult using *in vitro* or *in vivo* experimental systems. Hence, computational modelling provides a great opportunity to gain further mechanistic insight and to identify potential innovative drug targets and treatment strategies. Our simulation results using the genome sequence of SARS-CoV-2 identified the essential amino acid isoleucine as a rate-limiting metabolite. To our knowledge, such a dependency has not been reported in the literature. One study [60] linked lower isoleucine biosynthesis capability by the gut microbiome of severe/critical COVID-19 patients to higher inflammation. While this observation seems counterintuitive to our results, it is not known whether these patients had a more isoleucine-rich diet than healthy individuals, and thus a pre-infection microbiome low in isoleucine biosynthetic capabilities [60].

The viral load has been reported to be about ten times higher in patients with severe COVID-19 infections compared to the mild cases [54]. Consistently, for the WBM-virus models to remain feasible, we needed to increase the fraction of CD4+-T-cell, corresponding to T-cell activation, in the whole-body maintenance reaction to the reported increase of about 10 times (Table S3). This requirement is also consistent with our knowledge that immunocompromised individuals have a more severe COVID-19 infection outcome [2, 61, 62] and the central role of CD4+ T-cells in effective immune response and protection [63]. Thus, our host-virus model captures this aspect even though we do not explicitly model inflammation and immune response.

COVID-19 is primarily characterised by an infection of the respiratory tract [1] but its multisystemic nature has been well documented [4] and has been mainly attributed to the broad tissue expression of the hACE2 receptor [1]. In COVID-19 patients with severe infection or poor outcome, multi-organ failure has been observed [1], which raises the question of whether co-morbidities, such as liver damage, or the viral infection itself caused the multi-organ failures. Our computational modelling indicated a metabolic remodelling beyond the four organs that we infected *in silico* with SARS-CoV-2 (Figure 2, Table S4), while we did not model any organ damages. These results suggest that SARS-CoV-2, and likely any viral infection, initiates a metabolic remodelling on the whole-body level and not only on the level of the infected cells [14, 58, 59]. We observed such remodelling also in our mild SARS-CoV-2 models (Figure 2, Table S4), consistent with recent findings that also asymptomatic and mild infections can lead to, e.g., cognitive impairment [12] and long covid [13], which is characterised by a range of symptoms, such as fatigue, cognitive impairment, and shortness of breath [64]. This feature of COVID-19 raises the question of whether the observed and predicted, whole-body metabolic changes may become irreversible via, e.g., metabolic imprinting, thereby leading to chronic disease [65].

The blood metabolome represents a multi-facet readout of whole-body metabolic activity and environmental cues. Hence, we predicted potential changes in the blood metabolome upon SARS-CoV-2 infection, with and without T-cell activation, as an alternative read-out of whole-body metabolic remodelling (Table S5). Comparison with published plasma metabolomic data [55] from COVID-19 patients and healthy controls generally overlapped well with our predictions, thus, illustrating that the WBMs can be used, in principle, to predict plasma metabolome changes. Furthermore, numerous reported metabolites are either diet and/or microbiome associated. For instance, the tryptophane metabolism is well known to be influenced by the microbiome, which has also been shown to alter significantly in individuals infected with mild and severe COVID [66]. We are currently not accounting for microbial metabolic activity, but it would be a valuable extension of the current effort as it also permits elucidate host-microbiome co-metabolism [21]. Moreover, in our models, the availability of carnitine and its derivative was limited by the defined diet uptake, while carnitine and associated lipids have been reported to change with infection and disease severity [67]. Similarly, biotin [68] is increased in COVID patients. Biotin is either produced by the microbiome [69] or taken up with the diet. Similarly, one study [68] reported an increase in theophylline in COVID-2 patients. This metabolite is a drug, which is used to prevent and treat shortness of breath in asthma and COPD patients (Medline/Wiki – REF). Hence, medication needs to be recorded and corrected (as a confounding variable) when conducting metabolomic analysis, as it is well studied that medication directly influences the plasma metabolome [70, 71].

Almost 70 potential host drug targets and corresponding FDA-approved drugs have been suggested based on host-virus protein-protein interaction data [56]. We expected that of the few host proteins that mapped onto the genes included in the WBM model, which nonetheless covers the metabolic function of nearly 1,700 genes (Figure 1), at least some would alter the maximally possible virus shedding rate (Figure 3A, 3B). However, the inhibition of these proteins did not reduce the virus shedding rate. This result may be explained by how these drug targets were identified. These drug targets were chosen based on protein-protein interaction experiments [72] indicating that there is a physical interaction between the virus and host protein. However, interruption of this physical interaction cannot be modelled with our approach. It is also notable that only a very small fraction of these potential drug targets represents metabolic enzymes. In contrast, a substantial reduction in maximal possible virus shedding rate was achieved by inhibiting the guanylate kinase 1, which also can activate anti-viral prodrugs [73]. Interestingly, similar flux reduction was also achieved by reducing the lung uptake of either isoleucine or threonine. Such reduction in uptake rates may be achieved by reducing the blood concentration of the respective amino acids, e.g., through an altered diet.

No definite link between diet and the susceptibility to COVID-19 infection and the disease severity has been established yet [74]. Nonetheless, during the pandemic, the world health organisation (WHO) has been promoting healthy eating to boost the immune system and lower the risk of chronic diseases^1^, which are risk factors for a more severe outcome of COVID-19 [2]. Our simulation results suggest a link between diet and maximally possible virus shedding rate and even propose a sex-specific effect (Figure 3B-D). Our *in silico* results can be explained due to the mechanistic nature of our computational models and are a direct consequence of the varied amino acid content (especially of isoleucine) in the different *in silico* diets. This observation is exemplified by the finding of an essential amino acid (isoleucine) limiting the virus replication capacity, leading to the plausible hypothesis that inter-individual diet differences may contribute to the broad range of observed disease outcomes and clinical phenotypes. However, only integrating infection and clinical data with nutrition data will enable the validation of this hypothesis that we derived from our simulations. Future epidemiological studies, using infection, transmission and nutritional data across various countries may be able to shed a light on the role of diet habits in COVID-19 as will clinical trials involving nutritional components [74].

Most studies have focused on the significance of mutations in the spike (S) protein, as these have direct implications on how well the virus may be able to enter the cells and the effectiveness of the developed vaccines. While mutations in the other proteins may have structural and functional consequences, our simulation results suggest that there could also be a link to how fast the virus can replicate in the lung and that increases in virus replication, and thus virus shedding may be achieved by adapting better to host metabolism. We observed a striking anticorrelation between isoleucine requirement and the predicted virus shedding rate (Figure 5A). In contrast, the threonine requirement correlated with the predicted virus shedding rate for all variants, except for omicron (Figure 5B). Notably, while the abundance of these two amino acids did not alter substantially in the variant sequences (Figure 5C), the fact that they occurred in highly abundant proteins decreased/increased the amino acid’s requirement for viral replication. To our knowledge, this observation has not yet been reported in the literature and complements structural considerations of the virus mutations. Many factors influence whether a mutant becomes dominant and has a more severe outcome for infected individuals, including isolation, contract tracing, and lockdowns, which slow done transmission as well as the vaccination status of the population. Hence, our predicted virus shedding rates cannot easily be correlated with virus dominance. Also, the availability of genome sequences of variants on public servers, such as GISAID, is not suitable for correlating our predictions with a prominence of a variant due to biases in the selection of samples to be sequenced or differences in sequence capabilities of the different countries [75]. Nonetheless, it is remarkable that the delta variant, which has dominated most countries for many months in 2021 and caused many deaths worldwide, was predicted to have the highest maximal possible shedding rate while having the lowest isoleucine and highest threonine requirements. In contrast, the omicron variant, which has higher infectiousness but results overall in less severe COVID-19 cases [75], has a predicted virus shedding rate than the parental variant. Consistently, the sub-variant of omicron BA.2, which currently dominates the northern European countries has a similar predicted virus shedding rate as omicron BA.1. Our results highlight that in addition to the structural implications of virus mutations, one should also consider host metabolism implications. Importantly, these observations could lead to further novel treatment strategies for viral infections, including dietary restrictions on amino acid intake.

Numerous assumptions underlie our computational WBM-SARS-COV-2 models. One of the chief assumptions is the steady-state of the modelled metabolic network. As we consider, e.g., the virus replication potential for a day, we can assume that metabolism is at a steady-state, as biochemical reaction rates generally occur at a millisecond to seconds time scale [76]. A consequence of the steady-state assumption is that we cannot predict any concentration changes, except for those changes occurring across the system’s boundaries (e.g., virus shedding rate or blood metabolite accumulation/depletion rates). Equally, our modelling approach does not capture the dynamic changes occurring during infection but rather predicts a final feasible steady-state. Moreover, our WBM-SARS-COV-2 models only capture metabolism and thus can only inform about virus-host metabolic interactions, while viral infections and immune response are associated with substantial changes in the regulatory and signalling machinery [59]. Similarly, we do not consider the effect of fever on enzymatic rates or gene expression and also do not alter the blood supply to the different organs, which corresponded in our simulations to the resting state [21]. Moreover, we do not model decreased oxygen blood saturation that has been reported in severe COVID-19 cases [77, 78]. However, it has been estimated that virus shedding occurs already two to three days before symptoms appear [79], justifying our choice of not further parameterising the models with symptoms. Our models should therefore be seen as a model best suited for the early stages of the infection. Finally, we only performed our simulations using a representative reference man and woman [31]. More extensive simulations need to be carried out by parameterising the WBM-SARS-COV-2 models with data from, e.g., vulnerable populations (e.g., elderly or obese individuals), as well as ethnicity-specific parameters.

Despite these assumptions and limitations, we believe that our modelling approach provides valuable insights and strengths, such as the generation of novel hypotheses in a sex-specific, whole-body yet organ resolved manner during COVID-19 infection. These hypotheses, such as the possibility to reduce the virus replication rate by restricting isoleucine availability in the diet can be translated into clinical research, delivering thereby additional targets for intervention. Notably, the overall computational modelling paradigm could be extended to other viruses, such as influenza and human pathogens.

## Acknowledgement

We would like to thank Drs Cyrillic Thinnes, Johannes Hertel, and Bronson Weston for valuable discussions and feedback on various versions of the manuscript. We gratefully acknowledge the authors from the originating laboratories responsible for obtaining the specimens, as well as the submitting laboratories where the genome data were generated and shared via GISAID, on which this research is based (Table S10-S20).

## Footnotes

1 http://www.emro.who.int/nutrition/nutrition-infocus/nutrition-advice-for-adults-during-the-covid-19-outbreak.html

## References

1. Yuki K, Fujiogi M, Koutsogiannaki S: COVID-19 pathophysiology: A review. Clin Immunol 2020, 215:108427.

2. Wang B, Li R, Lu Z, Huang Y: Does comorbidity increase the risk of patients with COVID-19: evidence from meta-analysis. Aging 2020, 12.

3. Lu X, Zhang L, Du H, Zhang J, Li YY, Qu J, Zhang W, Wang Y, Bao S, Li Y et al: SARS-CoV-2 Infection in Children. New England Journal of Medicine 2020.

4. Gavriatopoulou M, Korompoki E, Fotiou D, Ntanasis-Stathopoulos I, Psaltopoulou T, Kastritis E, Terpos E, Dimopoulos MA: Organ-specific manifestations of COVID-19 infection. Clin Exp Med 2020, 20(4):493–506.

5. V’Kovski P, Kratzel A, Steiner S, Stalder H, Thiel V: Coronavirus biology and replication: implications for SARS-CoV-2. Nature reviews 2021, 19(3):155–170.

6. Amin M: COVID-19 and the liver: overview. Eur J Gastroenterol Hepatol 2021, 33(3):309–311.

7. Uhlen M, Oksvold P, Fagerberg L, Lundberg E, Jonasson K, Forsberg M, Zwahlen M, Kampf C, Wester K, Hober S et al: Towards a knowledge-based Human Protein Atlas. Nat Biotechnol 2010, 28(12):1248–1250.

8. Hamming I, Timens W, Bulthuis ML, Lely AT, Navis G, van Goor H: Tissue distribution of ACE2 protein, the functional receptor for SARS coronavirus. A first step in understanding SARS pathogenesis. J Pathol 2004, 203(2):631–637.

9. Lamers MM, Beumer J, van der Vaart J, Knoops K, Puschhof J, Breugem TI, Ravelli RBG, Paul van Schayck J, Mykytyn AZ, Duimel HQ et al: SARS-CoV-2 productively infects human gut enterocytes. Science (New York, NY 2020, 369(6499):50–54.

10. Wong SH, Lui RN, Sung JJ: Covid-19 and the Digestive System. J Gastroenterol Hepatol 2020.

11. Xiao F, Tang M, Zheng X, Liu Y, Li X, Shan H: Evidence for Gastrointestinal Infection of SARS-CoV-2. Gastroenterology 2020, 158(6):1831–1833 e1833.

12. Fernandez-Castaneda A, Lu P, Geraghty AC, Song E, Lee MH, Wood J, Yalcin B, Taylor KR, Dutton S, Acosta-Alvarez L et al: Mild respiratory SARS-CoV-2 infection can cause multi-lineage cellular dysregulation and myelin loss in the brain. bioRxiv 2022.

13. Taquet M, Geddes JR, Husain M, Luciano S, Harrison PJ: 6-month neurological and psychiatric outcomes in 236 379 survivors of COVID-19: a retrospective cohort study using electronic health records. Lancet Psychiatry 2021, 8(5):416–427.

14. Thaker SK, Ch’ng J, Christofk HR: Viral hijacking of cellular metabolism. BMC Biol 2019, 17(1):59.

15. Palsson BO: Systems Biology: Constraint-based Reconstruction and Analysis. UK: Cambridge university press; 2015.

16. Thiele I, Palsson BØ: A protocol for generating a high-quality genome-scale metabolic reconstruction. Nature protocols 2010, 5(1):93–121.

17. Mardinoglu A, Nielsen J: Systems medicine and metabolic modelling. J Intern Med 2012, 271(2):142–154.

18. Aurich MK, Thiele I: Computational Modeling of Human Metabolism and Its Application to Systems Biomedicine. Methods in molecular biology (Clifton, NJ 2016, 1386:253–281.

19. Bordbar A, Lewis NE, Schellenberger J, Palsson BO, Jamshidi N: Insight into human alveolar macrophage and M. tuberculosis interactions via metabolic reconstructions. Molecular systems biology 2010, 6:422.

20. Sahoo S, Thiele I: Predicting the impact of diet and enzymopathies on human small intestinal epithelial cells. Hum Mol Genet 2013, 22(13):2705–2722.

21. Thiele I, Sahoo S, Heinken A, Hertel J, Heirendt L, Aurich MK, Fleming RM: Personalized whole-body models integrate metabolism, physiology, and the gut microbiome. Molecular systems biology 2020, 16(5):e8982.

22. Shoaie S, Ghaffari P, Kovatcheva-Datchary P, Mardinoglu A, Sen P, Pujos-Guillot E, de Wouters T, Juste C, Rizkalla S, Chilloux J et al: Quantifying Diet-Induced Metabolic Changes of the Human Gut Microbiome. Cell metabolism 2015, 22(2):320–331.

23. Sahoo S, Franzson L, Jonsson JJ, Thiele I: A compendium of inborn errors of metabolism mapped onto the human metabolic network. Molecular bioSystems 2012, 8(10):2545–2558.

24. Cheng Y, Schlosser P, Hertel J, Sekula P, Oefner PJ, Spiekerkoetter U, Mielke J, Freitag DF, Schmidts M, Investigators G et al: Rare genetic variants affecting urine metabolite levels link population variation to inborn errors of metabolism. Nat Commun 2021, 12(1):964.

25. Hyduke DR, Lewis NE, Palsson BO: Analysis of omics data with genome-scale models of metabolism. Molecular bioSystems 2013, 9(2):167–174.

26. Saha R, Chowdhury A, Maranas CD: Recent advances in the reconstruction of metabolic models and integration of omics data. Curr Opin Biotechnol 2014, 29:39–45.

27. Preciat Gonzalez GA: XomicsToModel: Multiomics data integration and generation of thermodynamically consistent metabolic models. bioRxiv 2021.

28. Renz A, Widerspick L, Drager A: FBA reveals guanylate kinase as a potential target for antiviral therapies against SARS-CoV-2. Bioinformatics (Oxford, England) 2020, 36(Suppl 2):i813–i821.

29. Renz A, Widerspick L, Drager A: Genome-Scale Metabolic Model of Infection with SARS-CoV-2 Mutants Confirms Guanylate Kinase as Robust Potential Antiviral Target. Genes (Basel) 2021, 12(6).

30. Cheng K, Martin-Sancho L, Pal LR, Pu Y, Riva L, Yin X, Sinha S, Nair NU, Chanda SK, Ruppin E: Genome-scale metabolic modeling reveals SARS-CoV-2-induced metabolic changes and antiviral targets. Molecular systems biology 2021, 17(11):e10260.

31. Snyder WS, Cook MJ, Karhausen LR, Nasset ES, Parry Howells G, Tipton IH: Report on the Task Group on Reference Man; 1975.

32. Brunk E, Sahoo S, Zielinski DC, Altunkaya A, Drager A, Mih N, Gatto F, Nilsson A, Preciat Gonzalez GA, Aurich MK et al: Recon3D enables a three-dimensional view of gene variation in human metabolism. Nat Biotechnol 2018, 36(3):272–281.

33. Wishart DS, Jewison T, Guo AC, Wilson M, Knox C, Liu Y, Djoumbou Y, Mandal R, Aziat F, Dong E et al: HMDB 3.0--The Human Metabolome Database in 2013. Nucleic Acids Res 2013, 41(Database issue):D801–807.

34. Thiele I, Fleming RM, Bordbar A, Schellenberger J, Palsson BO: Functional characterization of alternate optimal solutions of Escherichia coli’s transcriptional and translational machinery. Biophysical journal 2010, 98(10):2072–2081.

35. Fukushima H, Hoshina K, Nakamura R, Ito Y, Gomyoda M: Epidemiological study of Yersinia enterocolitica and Yersinia pseudotuberculosis in Shimane Prefecture, Japan. Contrib Microbiol Immunol 1987, 9:103–110.

36. Klumperman J, Locker JK, Meijer A, Horzinek MC, Geuze HJ, Rottier PJ: Coronavirus M proteins accumulate in the Golgi complex beyond the site of virion budding. Journal of virology 1994, 68(10):6523–6534.

37. Neuman BW, Adair BD, Yoshioka C, Quispe JD, Orca G, Kuhn P, Milligan RA, Yeager M, Buchmeier MJ: Supramolecular architecture of severe acute respiratory syndrome coronavirus revealed by electron cryomicroscopy. Journal of virology 2006, 80(16):7918–7928.

38. Sturman LS, Holmes KV, Behnke J: Isolation of coronavirus envelope glycoproteins and interaction with the viral nucleocapsid. Journal of virology 1980, 33(1):449–462.

39. Godet M, L’Haridon R, Vautherot JF, Laude H: TGEV corona virus ORF4 encodes a membrane protein that is incorporated into virions. Virology 1992, 188(2):666–675.

40. Neuman BW, Joseph JS, Saikatendu KS, Serrano P, Chatterjee A, Johnson MA, Liao L, Klaus JP, Yates JR, 3rd, Wuthrich K et al: Proteomics analysis unravels the functional repertoire of coronavirus nonstructural protein 3. Journal of virology 2008, 82(11):5279–5294.

41. Wrapp D, Wang N, Corbett KS, Goldsmith JA, Hsieh CL, Abiona O, Graham BS, McLellan JS: Cryo-EM structure of the 2019-nCoV spike in the prefusion conformation. Science (New York, NY 2020, 367(6483):1260–1263.

42. Fung TS, Liu DX: Post-translational modifications of coronavirus proteins: roles and function. Future Virol 2018, 13(6):405–430.

43. McBride CE, Machamer CE: Palmitoylation of SARS-CoV S protein is necessary for partitioning into detergent-resistant membranes and cell-cell fusion but not interaction with M protein. Virology 2010, 405(1):139–148.

44. Guan X, Fierke CA: Understanding Protein Palmitoylation: Biological Significance and Enzymology. Sci China Chem 2011, 54(12):1888–1897.

45. Aller S, Scott A, Sarkar-Tyson M, Soyer OS: Integrated human-virus metabolic stoichiometric modelling predicts host-based antiviral targets against Chikungunya, Dengue and Zika viruses. J R Soc Interface 2018, 15(146).

46. Bar-On YM, Flamholz A, Phillips R, Milo R: SARS-CoV-2 (COVID-19) by the numbers. eLife 2020, 9.

47. Vankadari N, Wilce JA: Emerging WuHan (COVID-19) coronavirus: glycan shield and structure prediction of spike glycoprotein and its interaction with human CD26. Emerg Microbes Infect 2020, 9(1):601–604.

48. Thevarajan I, Nguyen THO, Koutsakos M, Druce J, Caly L, van de Sandt CE, Jia X, Nicholson S, Catton M, Cowie B et al: Breadth of concomitant immune responses prior to patient recovery: a case report of non-severe COVID-19. Nature Medicine 2020.

49. Noronha A, Modamio J, Jarosz Y, Guerard E, Sompairac N, Preciat G, Danielsdottir AD, Krecke M, Merten D, Haraldsdottir HS et al: The Virtual Metabolic Human database: integrating human and gut microbiome metabolism with nutrition and disease. Nucleic Acids Res 2019, 47(D1):D614–D624.

50. Orth JD, Thiele I, Palsson BO: What is flux balance analysis? Nat Biotechnol 2010, 28(3):245–248.

51. Noronha A, Danielsdottir AD, Gawron P, Johannsson F, Jonsdottir S, Jarlsson S, Gunnarsson JP, Brynjolfsson S, Schneider R, Thiele I et al: ReconMap: an interactive visualization of human metabolism. Bioinformatics (Oxford, England) 2017, 33(4):605–607.

52. Heirendt L, Arreckx S, Pfau T, Mendoza SN, Richelle A, Heinken A, Haraldsdottir HS, Wachowiak J, Keating SM, Vlasov V et al: Creation and analysis of biochemical constraint-based models using the COBRA Toolbox v.3.0. Nature protocols 2019, 14(3):639–702.

53. Buchfink B, Xie C, Huson DH: Fast and sensitive protein alignment using DIAMOND. Nature methods 2015, 12(1):59–60.

54. Chen Y, Li L: SARS-CoV-2: virus dynamics and host response. Lancet Infect Dis 2020.

55. Shen B, Yi X, Sun Y, Bi X, Du J, Zhang C, Quan S, Zhang F, Sun R, Qian L et al: Proteomic and Metabolomic Characterization of COVID-19 Patient Sera. Cell 2020, 182(1):59–72 e15.

56. Gordon DE, Jang GM, Bouhaddou M, Xu J, Obernier K, O’Meara MJ, Guo JZ, Swaney DL, Tummino TA, Huettenhain R et al: A SARS-CoV-2-Human Protein-Protein Interaction Map Reveals Drug Targets and Potential Drug-Repurposing. bioRxiv 2020:2020.2003.2022.002386.

57. Colson P, Delerce J, Burel E, Dahan J, Jouffret A, Fenollar F, Yahi N, Fantini J, La Scola B, Raoult D: Emergence in southern France of a new SARS-CoV-2 variant harbouring both N501Y and E484K substitutions in the spike protein. Arch Virol 2022, 167(4):1185–1190.

58. Goodwin CM, Xu S, Munger J: Stealing the Keys to the Kitchen: Viral Manipulation of the Host Cell Metabolic Network. Trends Microbiol 2015, 23(12):789–798.

59. Raniga K, Liang C: Interferons: Reprogramming the Metabolic Network against Viral Infection. Viruses 2018, 10(1).

60. Zhang F, Wan Y, Zuo T, Yeoh YK, Liu Q, Zhang L, Zhan H, Lu W, Xu W, Lui GCY et al: Prolonged Impairment of Short-Chain Fatty Acid and L-Isoleucine Biosynthesis in Gut Microbiome in Patients With COVID-19. Gastroenterology 2022, 162(2):548–561 e544.

61. Petrilli CM, Jones SA, Yang J, Rajagopalan H, O’Donnell LF, Chernyak Y, Tobin K, Cerfolio RJ, Francois F, Horwitz LI: Factors associated with hospitalization and critical illness among 4,103 patients with COVID-19 disease in New York City. medRxiv 2020:2020.2004.2008.20057794.

62. Drucker DJ: Coronavirus infections and type 2 diabetes-shared pathways with therapeutic implications. Endocrine reviews 2020:bnaa011.

63. Swain SL, McKinstry KK, Strutt TM: Expanding roles for CD4(+) T cells in immunity to viruses. Nature reviews 2012, 12(2):136–148.

64. Crook H, Raza S, Nowell J, Young M, Edison P: Long covid-mechanisms, risk factors, and management. Bmj 2021, 374:n1648.

65. Waterland RA, Garza C: Potential mechanisms of metabolic imprinting that lead to chronic disease. Am J Clin Nutr 1999, 69(2):179–197.

66. Yamamoto S, Saito M, Tamura A, Prawisuda D, Mizutani T, Yotsuyanagi H: The human microbiome and COVID-19: A systematic review. PLoS One 2021, 16(6):e0253293.

67. Li C, Ou R, Wei Q, Shang H: Carnitine and COVID-19 Susceptibility and Severity: A Mendelian Randomization Study. Front Nutr 2021, 8:780205.

68. Blasco H, Bessy C, Plantier L, Lefevre A, Piver E, Bernard L, Marlet J, Stefic K, Benz-de Bretagne I, Cannet P et al: The specific metabolome profiling of patients infected by SARS-COV-2 supports the key role of tryptophan-nicotinamide pathway and cytosine metabolism. Scientific reports 2020, 10(1):16824.

69. Magnusdottir S, Ravcheev D, de Crecy-Lagard V, Thiele I: Systematic genome assessment of B-vitamin biosynthesis suggests co-operation among gut microbes. Front Genet 2015, 6:148.

70. Krauss RM, Zhu H, Kaddurah-Daouk R: Pharmacometabolomics of statin response. Clin Pharmacol Ther 2013, 94(5):562–565.

71. Elbadawi-Sidhu M, Baillie RA, Zhu H, Chen YI, Goodarzi MO, Rotter JI, Krauss RM, Fiehn O, Kaddurah-Daouk R: Pharmacometabolomic signature links simvastatin therapy and insulin resistance. Metabolomics : Official journal of the Metabolomic Society 2017, 13.

72. !!! INVALID CITATION !!! [76].

73. Hible G, Daalova P, Gilles AM, Cherfils J: Crystal structures of GMP kinase in complex with ganciclovir monophosphate and Ap5G. Biochimie 2006, 88(9):1157–1164.

74. James PT, Ali Z, Armitage AE, Bonell A, Cerami C, Drakesmith H, Jobe M, Jones KS, Liew Z, Moore SE et al: The Role of Nutrition in COVID-19 Susceptibility and Severity of Disease: A Systematic Review. J Nutr 2021, 151(7):1854–1878.

75. Maxmen A: Omicron blindspots: why it’s hard to track coronavirus variants. Nature 2021, 600(7890):579.

76. Papin JA, Hunter T, Palsson BO, Subramaniam S: Reconstruction of cellular signalling networks and analysis of their properties. Nat Rev Mol Cell Biol 2005, 6(2):99–111.

77. Yang W, Cao Q, Qin L, Wang X, Cheng Z, Pan A, Dai J, Sun Q, Zhao F, Qu J et al: Clinical characteristics and imaging manifestations of the 2019 novel coronavirus disease (COVID-19):A multi-center study in Wenzhou city, Zhejiang, China. J Infect 2020, 80(4):388–393.

78. Grasselli G, Zangrillo A, Zanella A, Antonelli M, Cabrini L, Castelli A, Cereda D, Coluccello A, Foti G, Fumagalli R et al: Baseline Characteristics and Outcomes of 1591 Patients Infected With SARS-CoV-2 Admitted to ICUs of the Lombardy Region, Italy. JAMA 2020.

79. He X, Lau EHY, Wu P, Deng X, Wang J, Hao X, Lau YC, Wong JY, Guan Y, Tan X et al: Temporal dynamics in viral shedding and transmissibility of COVID-19. Nature Medicine 2020.

